# A quantitative framework for evaluating single-cell data structure preservation by dimensionality reduction techniques

**DOI:** 10.1101/684340

**Authors:** Cody N. Heiser, Ken S. Lau

**Affiliations:** Epithelial Biology Center, Vanderbilt University Medical Center, 2213 Garland Avenue, 10475 MRB IV, Nashville, TN 37232, USA; Program in Chemical and Physical Biology, Vanderbilt University School of Medicine, Nashville, TN 37232, USA; Department of Cell and Developmental Biology, Vanderbilt University School of Medicine, Nashville, TN 37232, USA; Center for Quantitative Sciences, Vanderbilt University School of Medicine, Nashville, TN 37232, USA

**Keywords:** Dimensionality reduction, single-cell transcriptomics, single-cell analysis, unsupervised learning

## Abstract

High-dimensional data, such as those generated using single-cell RNA sequencing, present challenges in interpretation and visualization. Numerical and computational methods for dimensionality reduction allow for low-dimensional representation of genome-scale expression data for downstream clustering, trajectory reconstruction, and biological interpretation. However, a comprehensive and quantitative evaluation of the performance of these techniques has not been established. We present an unbiased framework that defines metrics of global and local structure preservation in dimensionality reduction transformations. Using discrete and continuous scRNA-seq datasets, we find that input cell distribution and method parameters are largely determinant of global, local, and organizational data structure preservation by eleven published dimensionality reduction methods. Code available at github.com/KenLauLab/DR-structure-preservation allows for rapid evaluation of further datasets and methods.

## Introduction

Single-cell RNA sequencing (scRNA-seq) offers parallel, genome-scale measurement of tens of thousands of transcripts for thousands of cells (Klein *et al*., 2015; Macosko *et al*., 2015). Data of this magnitude provide powerful insight toward cell identity and developmental trajectory – states and fates – which are used to interrogate tissue heterogeneity and characterize disease progression (Regev *et al*., 2017; Wagner *et al*., 2019). Yet, extracting meaningful information from such high-dimensional data presents a massive challenge. Numerical and computational methods for dimensionality reduction have been developed to reconstruct underlying distributions from native “gene space” and provide low-dimensional, latent representations of single-cell data for more intuitive downstream interpretation. Basic clustering methods and linear transformations such as principal component analysis (PCA) have proven to be valuable tools in this field (Sorzano, Vargas and Montano, 2014; Levine *et al*., 2015; Kiselev *et al*., 2017; Tsuyuzaki *et al*., 2019). However, given the distribution and sparsity of scRNA-seq data, complex, nonlinear transformations are often required to capture and visualize expression patterns. Unsupervised machine learning techniques and, more recently, deep learning methods, are being rapidly developed to assist researchers in single-cell transcriptomic analysis (Van der Maaten and Hinton, 2008; Pierson and Yau, 2015; Linderman *et al*., 2017; Wang *et al*., 2017; Becht *et al*., 2018; Ding, Condon and Shah, 2018; Lopez *et al*., 2018; Risso *et al*., 2018; Eraslan *et al*., 2019; Townes *et al*., 2019). Because these techniques condense cell features in the native space to a small number of latent dimensions for visualization, lost information can result in exaggerated or dampened cell-cell similarity. Furthermore, depending on input data and user-defined parameters, the structure of resulting embeddings can vary greatly, potentially altering biological interpretation (Kobak and Berens, 2019).

With a deluge of computational techniques for dimensionality reduction, the field is lacking a comprehensive assessment of native organizational distortion consequential to such methods. Characterization of these tools would enable end users to confidently determine the most suitable methods for their application. We present an unbiased, quantitative framework for evaluation of data structure preservation by dimensionality reduction transformations. We propose three metrics for broad characterization of strengths and weaknesses of these methods based on cell-cell distance in native gene space. Initial benchmarking of eleven published software tools on discrete and continuous cell distributions shows global, local, and organizational data structure conservation under different parameter and input conditions. Interpretation and best practices, as well as extensibility of this method to other dimensionality reduction tools, is discussed.

## Results

### Cell distance distributions describe global structure of high-dimensional datasets

In order to evaluate dimensionality reduction techniques, Euclidean cell-cell distance in native, high-dimensional space is used as a quantitative standard. Counts of unique molecular identifiers (UMI) for each gene make up the features, or columns of the dataset, while every observation, or row, represents a single cell (Figure 1A). In this way, transcriptomic data can be represented as an *m × n* matrix (cells x genes).

**Figure 1.**
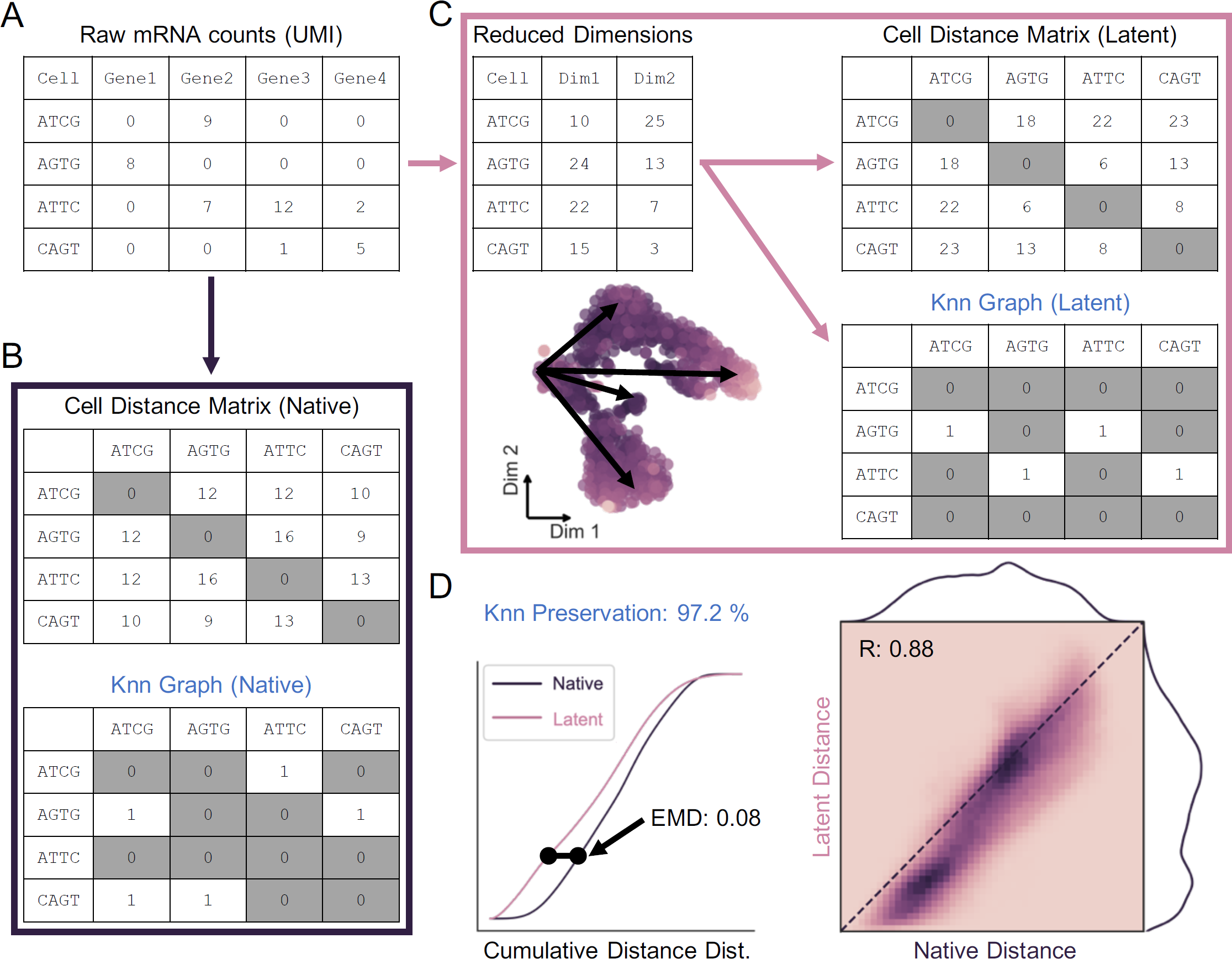
Cell distance distributions describe global structure of high-dimensional datasets. A) Representation of gene counts matrix as output by scRNA-seq processing pipeline. B) Cell-cell Euclidean distances in native gene space are calculated to generate an m *×* m matrix, where m is the total number of cells. K nearest-neighbors (Knn) graph is constructed from these distances as a binary m *×* m matrix. C) Upon transformation to low-dimensional latent space, a distance matrix and Knn graph can be calculated as in B. D) Distance matrices from native (B) and latent (C) spaces are used to build cumulative probability density distributions, which can be compared to one another by Earth Mover’s Distance (EMD, left). Unique distances from the upper triangle of each distance matrix are correlated by Mantel test (right). Knn preservation represents element-wise comparison of nearest-neighbor graph matrices in each space. See Methods.

The global data structure in the native space can be constructed by first calculating an *m × m* matrix that contains the pairwise distances between all observations in *n* dimensions (Figure 1B, top). The upper triangle of this distance matrix contains all unique cell distances in the dataset, which can then be represented by a probability density distribution as in Figure 1D. From these distances, local “neighborhoods” can be defined in the form of a K nearest-neighbor (Knn) graph. The Knn graph is represented by a binary *m × m* matrix that defines the K cells with the shortest distances from each cell in the dataset (Figure 1B, bottom). Similarly, a distance matrix, distance distribution, and Knn graph can be constructed from a low-dimensional latent space resulting from dimensionality reduction (Figure 1C).

Overall distance preservation following dimensionality reduction is measured by Mantel correlation (Figure 1D, right). This method was designed for symmetrical matrices that represent element-wise similarity between two vectors, and appropriately accounts for multiple testing of 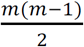 distances (Mantel, 1967). Structural alteration of the cell distance distribution – constructed from the upper triangle of the distance matrix – is quantified by the Wasserstein metric, or Earth-Mover’s Distance (EMD) (Figure 1D, left). Widely applied to image processing, this metric determines the energy associated with shifting one distribution to another, and can also be defined as the area between two cumulative probability distributions (Werman, Peleg and Rosenfeld, 1985; Rubner, Tomasi and Guibas, 1998, 2000; Levina and Bickel, 2001). Finally, preservation of the Knn graph before and after low-dimensional embedding can also be quantified as the percentage of total binary matrix elements conserved in order to describe maintenance of local substructures in the data.

### Discrete and continuous cell distributions exemplify common biological patterns *in vivo*

A major consideration for testing dimensionality reduction techniques is the true structure of the input data in native, high-dimensional space. For the scope of our evaluation applied to single-cell transcriptomics, we identify two overarching classes of scRNA-seq data for proof-of-principal: discrete and continuous. Discrete single-cell data are comprised of differentiated cell types with unique, highly discernable gene expression profiles. These data include classic PBMC experiments and neuronal datasets which can be easily clustered into distinct cell types (Zeisel *et al*., 2015; Rheaume *et al*., 2018). Conversely, continuous data represent multi-faceted expression gradients present during cell development and differentiation, and are commonly associated with dynamic systems such as erythropoiesis or embryonic development (Tusi *et al*., 2018; Wagner *et al*., 2019). Recently, computational tools for trajectory inference and lineage reconstruction from these data are being rapidly developed to query differential expression and gene-regulatory events involved in cell fate decisions (Qiu *et al*., 2017; Herring, Chen, *et al*., 2018; Saelens *et al*., 2019; Van den Berge *et al*., 2019).

Mouse retina cells, analyzed using Drop-seq by Macosko and coworkers, provide a discrete cell distribution for our analysis (Macosko *et al*., 2015) (GEO accession ID GSM1626793). Counts data from 20,478 genes for 1,326 cells were analyzed using PhenoGraph to determine “ground-truth” cell clusters by the Louvain algorithm (Figure 2A) (Levine *et al*., 2015). We performed relatively coarse clustering, ignoring subtype heterogeneity in favor of clusters reflecting principal cell identity amenable to our downstream analyses (see Methods). A t-SNE projection primed with 100 principal components (PCs) of all transcript counts allows for visualization of the data structure and represented cell types (Figure 2B). As evident from the 2D embedding, these data are highly discrete, and constituent cell clusters are easily distinguished by gene expression (Figure 2C, Figure S2A).

**Figure 2.**
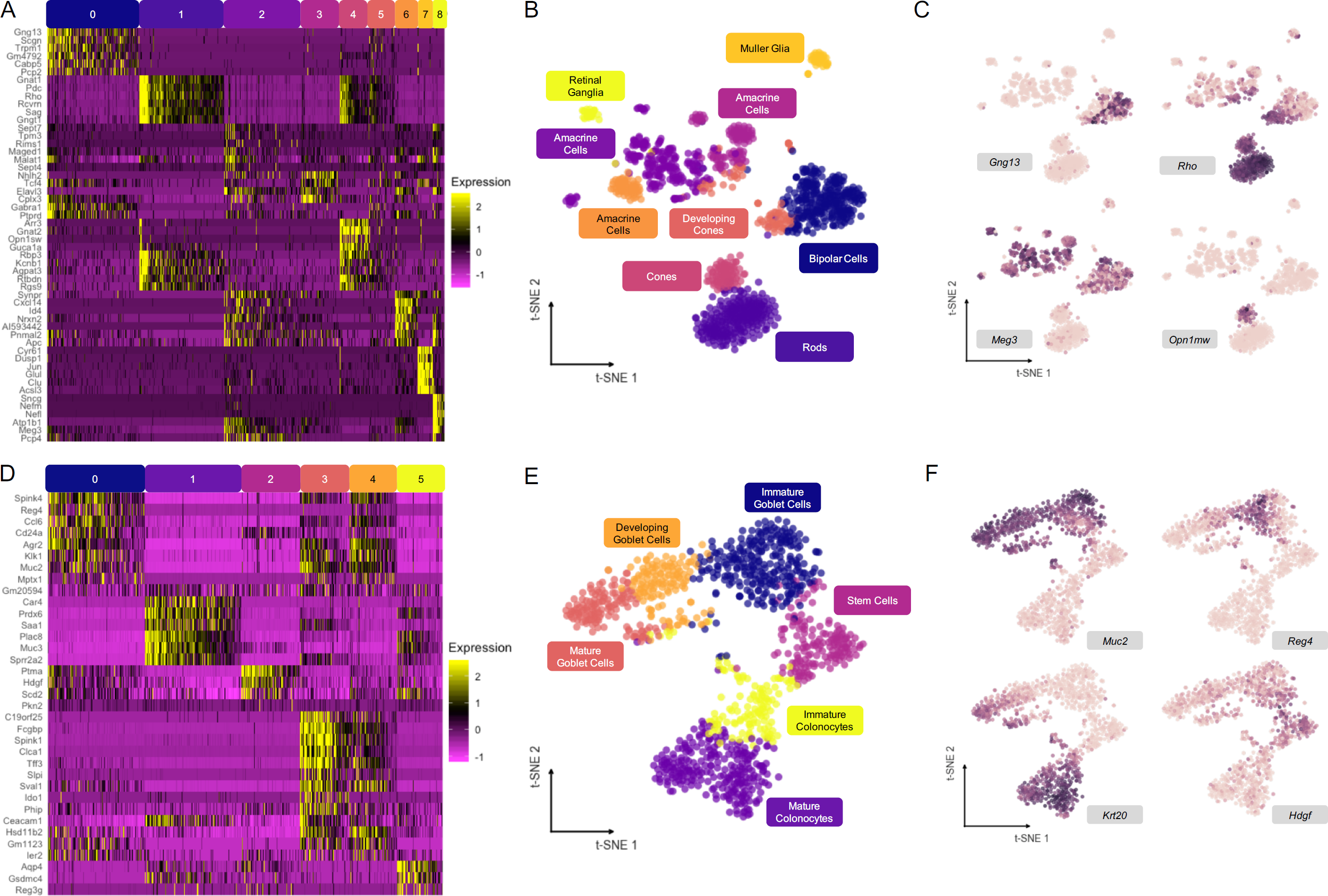
Discrete and continuous cell distributions exemplify common biological patterns *in vivo*. A) Relative expression of top variable genes in each Louvain cluster for mouse retina dataset (Macosko *et al*., 2015). B) t-SNE embedding primed with 100 principal components (PCs) of retina dataset with overlay of consensus clusters. C) t-SNE projection from B with overlay of example marker genes used to identify cell types in A. Expression represented as arcsinh-normalized raw counts. D-F) Same as in A-C, for mouse colonic epithelium dataset (Herring *et al*., 2018). See Methods.

Mouse colon data, representing a continuous distribution of actively differentiating cells along the crypt-villus axis of the colonic epithelium, were generated using inDrops scRNA-seq (Herring, Banerjee, *et al*., 2018) (GEO accession ID GSM2743164). Counts data from 25,504 genes for 1,117 cells were similarly analyzed by PhenoGraph and t-SNE to visualize continuous data structure (Figure 2D,E). The six clusters form a branching continuum of cell states identified by expression markers (Figure 2F, Figure S2B), resolving two major lineages in the colon: absorptive colonocytes and secretory goblet cells (Lepourcelet *et al*., 2005; Tamura *et al*., 2007; Larsson *et al*., 2012). These clusters are linked together by pseudotemporal trajectories and thus their arrangement is expected to be conserved upon low-dimensional embedding into a latent space.

### Input cell distribution determines performance of global structure preservation

Using the metrics outlined in Figure 1, we compared eleven published dimensionality reduction techniques applied to continuous and discrete datasets. To allow for direct input to dimensionality reduction tools, raw counts for both datasets were feature-selected to the 500 most variable genes. This “brute force” feature selection technique may not be optimal for enriching rare cell types for higher-level analyses (Chen, Herring and Lau, 2018). However, for our purpose of proof-of-principal valuation of dimensionality reduction methods, it easily provides a high level of differential information for unsupervised algorithms without biasing data input. Calculating our metrics on all cells in the dataset, we first assess global structure preservation following transformation to a dimension-reduced latent space. Representative examples of two-dimensional projections and their corresponding distance distributions and correlations using SIMLR for the retina dataset and UMAP for the colon dataset are shown in Figure 3A and Figure 3E, respectively. Notably, the largest discrepancy in structural preservation is between the two datasets, highlighting the significance of input cell distribution to overall method performance. Intuitively, Knn preservation is higher for all eleven methods when applied to the colon dataset, reflecting the notion of continuous neighborhoods – a moving window of expression gradients – connecting all cells through developmental pseudotime. Another important observation regarding the dimension-reduced latent spaces involves the directionality of the cell distance distribution shift. A compression of distances from native to latent space is indicated by a shift left in the cumulative distance distribution (Figure 3B,F, Figure S1A) or below the identity line in the unique distance correlation (Figure 3D,H, Figure S1B). Alternatively, a shift right in the cumulative distance distribution or above the identity line of the distance correlation signifies an exaggeration of native distances (Figure S1). These phenomena are important in the context of global versus local structure preservation. For example, UMAP appears to compress small, local distances to a greater extent than t-SNE, while both methods maintain relative global structure as indicated by a favorable correlation of large distances between clusters. Although this characteristic of UMAP embeddings causes greater information loss reflected in less favorable preservation metrics (Figure 3C,G), clusters within the resulting projections tend to be highly condensed and perhaps more easily interpreted (Figure S3A,B).

**Figure 3.**
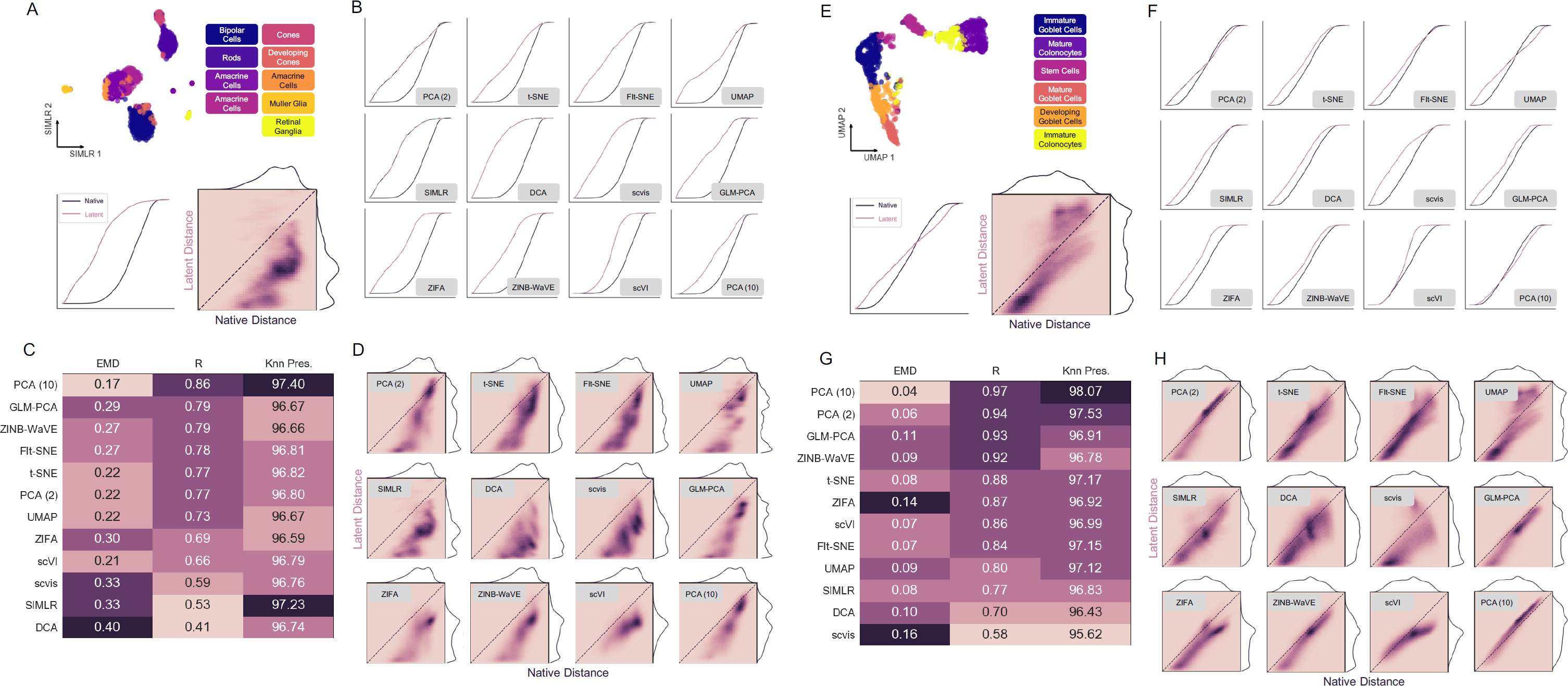
Global structure preservation analysis of 11 dimensionality reduction methods on discrete and continuous scRNA-seq datasets. A) Example 2D projection of mouse retina data using SIMLR with Louvain cluster overlay (top). Cumulative distance distributions for native and latent spaces (bottom left) and 2D histogram representing Mantel correlation between the two (bottom right). B) Cumulative distance distributions of evaluated latent spaces for mouse retina data. C) Summary of structure preservation metrics for 11 dimensionality reduction methods on mouse retina dataset. D) 2D histograms of cell distance correlations of evaluated latent spaces for mouse retina data. E) Same as in A, with UMAP projection of mouse colon data. F-H) Same as in B-D, for mouse colon data. 10-component PCA is shown for direct comparison to scVI’s 10-dimensional latent space; all other methods return 2D embedding. See Methods.

### Parameter optimization plays key role in structural preservation

User-defined parameters for unsupervised algorithms often present themselves as “black-box” knobs with unknown consequences. Tuning these parameters can be a daunting task for the single-cell analyst, but is known to be crucial to algorithm performance (Belkina *et al*., 2018; Kobak and Berens, 2019; Tsuyuzaki *et al*., 2019). Using our proposed metrics, we evaluated global structure preservation across a range of perplexity values for t-SNE and UMAP algorithms applied to both discrete and continuous data. Through a balance of distance correlation, EMD, and Knn preservation, we can identify an initial range of optimal perplexity values between 2 and 10 % of the total number of cells in the dataset (Figure S3C).

### Substructure analysis illuminates contribution to global performance

To corroborate results of global structure preservation and dissect contribution of local (within cluster) and organizational (between cluster) distances to overall dimensionality reduction performance, clusters were isolated for targeted substructure quantification. Here, we can measure distance preservation of individual clusters as well as distances between clusters to emphasize local arrangement (Figure S1C,D).

Retinal cone cells (Figure 2A, cluster 4, *n* = 94) were used as an example of local distances in the discrete dataset, while mature colonocytes (Figure 2D, cluster 1, *n* = 273) were isolated in the colon dataset (Figure 4A,B,E,F). Local distance compression represents the overarching trend for the eleven evaluated tools, indicated by a correlation shift below the identity line. The latent spaces from scVI and 10-component PCA are notable exceptions, yielding the two lowest EMD values for each dataset (Figure S4A). This most likely results from the 10-dimensional latent spaces of these methods capturing more cellular variability than 2D projections. Added noise in the SIMLR latent space of mouse retina cells indicates a disagreement with Louvain cluster membership, and may be attributed to the truncated, 500-feature input used for our analysis (Figure 4B). Moreover, this observation suggests that discrete, “on-off” expression patterns are less robust to dropouts that cause misassignment of cell type than continuous gradients of gene expression.

Besides maintenance of intra-cluster local structure, dimensionality reduction methods are also tasked with preserving cellular neighborhoods, or relationships between clusters. By calculating distance distributions from cells in one cluster to those in another, we can evaluate these associations. Furthermore, we can analyze pairwise cluster-cluster distances to investigate organization of data substructures (Figure S1C). In the mouse retina dataset, distances between bipolar cells, rod cells, and amacrine cells (Figure 2A, clusters 0, 1, 2, *n* = 309, 281, 258) are marked largely by compression, with some tools altering the arrangement of the three clusters (Figure 4D, red boxes). For example, the bipolar and amacrine clusters are closest to one another in the native gene space, but the bipolar cell cluster is closer to the rod cell cluster in the UMAP embedding, as indicated by the ordering of each distribution on the axes of the 2D histogram plot. Conversely, relative distances between three adjacent clusters along the goblet cell lineage (Figure 2D, clusters 0, 3 and 4, *n* = 274, 140 and 135) are more highly conserved by all dimensionality reductions. These results confirm that related cells in continuous scRNA-seq data are tethered to their neighbors through intermediate expression states, resulting in improved local structure preservation upon latent projection (Figure S4).

## Discussion

Single-cell RNA sequencing (scRNA-seq) allows for high-throughput, genome-scale measurement of mRNA expression in individual cells. Interpretation, pattern detection, and visualization of such high-dimensional observations present major challenges, and current datasets are constantly expanding in breadth and resolution. Systems biologists have derived and adapted numerical and computational methods for dimensionality reduction to allow for low-dimensional representation of single-cell data and deduction of cell states and fates (Van der Maaten and Hinton, 2008; Pierson and Yau, 2015; Linderman *et al*., 2017; Wang *et al*., 2017; Becht *et al*., 2018; Ding, Condon and Shah, 2018; Lopez *et al*., 2018; Risso *et al*., 2018; Eraslan *et al*., 2019; Townes *et al*., 2019). As software tools for single-cell analysis become widely available to lab-based researchers, there is a need to thoroughly understand how underlying biological information is maintained or distorted by these techniques. Two scRNA-seq datasets, representing discrete, differentiated cell types (Macosko *et al*., 2015) and continuous, hierarchical differentiation states (Herring, Banerjee, *et al*., 2018) were used to investigate cell distance preservation by eleven published dimensionality reduction methods. These dichotomous data offer insight into strengths and weaknesses of these tools for different applications. Distance correlation, EMD between distance distributions, and nearest-neighbor preservation were assessed to quantify cell dispersion, global data structure preservation, and neighborhood maintenance.

We identified dispersion trends in local and global distance distributions that denote expansion and contraction of native cell distances (Figure S1). This allowed us to evaluate general performance of dimensionality reduction methods on entire single-cell datasets (Figure 3), and take a deeper dive to examine how local distances – within or between clusters – contribute to the global structure of a low-dimensional latent space (Figure 4). With a goal of grouping cells by their gene expression profiles, most dimensionality reduction tools evaluated herein compressed local distances, embellishing cluster similarity, while maintaining or expanding global distances, exaggerating cluster distinction (Figure 3, Figure 4). These characteristics of dimensionality reduction methods are desirable for most applications. However, resolution of rare cell types and sub-cluster heterogeneity may be lost, stressing the importance of input data quality, feature selection, and user-defined parameters.

Discrete scRNA-seq data are more susceptible to structural perturbation by downstream dimensionality reduction, as indicated by larger EMD values and lower distance correlations in the retina dataset than colonic epithelial cells (Figure 3, Figure S4). The possibility of cell type misclassification, or “noisy” cluster membership in low-dimensional embeddings compared to consensus Louvain clustering, is exacerbated in discrete data. This noise is intensified by gene dropouts, and is therefore sensitive to sequencing depth and capture efficiency. We also observed cluster rearrangement within the retina dataset, suggesting that relative substructure organization is poorly defined for discrete datasets (Figure 4D, Figure S4). On the other hand, continuous cell distributions are more robust to these effects. In a continuum of gene expression, as in the actively differentiating cells along the crypt-villus axis of the colonic epithelium, cell clusters are tethered to one another through intermediate states. In this way, preservation of local and relative substructures is built-in to dimensionality reduction analysis of continuous datasets, and results in more credible representations of native data structure (Figure 4F,H, Figure S4). Finally, cursory exploration of the perplexity parameter in t-SNE and UMAP reveals a range of optimal values that yield favorable structure preservation metrics, endorsing the need for parameter optimization for dimensionality reduction of scRNA-seq datasets (Figure S3C).

As high-dimensional datasets become increasingly pervasive in systems biology, computational tools for reliable and reproducible analysis of these data become tremendous assets to discovery. Dimensionality reduction techniques allow for embedding cellular observations with tens of thousands of gene features into a low-dimensional space for visualization and downstream processing. Many such methods exist, calling on concepts from mathematics and computer science to aggregate underlying biological patterns into a latent representation of the data. We present an unbiased, quantitative framework based on native cell distance to evaluate data structure preservation by dimensionality reduction tools. All code associated with this project is available at github.com/KenLauLab/DR-structure-preservation, and is readily extensible to additional scRNA-seq datasets and dimensionality reduction methods.

## Acknowledgments

The authors would like to acknowledge the Vanderbilt Epithelial Biology Center and the Quantitative Systems Biology Center for helpful discussions. CNH and KSL are funded by NIH grants R01DK103831, U2CCA233291, and R01CA238553.

## Author Contributions

CNH and KSL conceived of the study. CNH developed methodology, analyzed the data, and generated visualizations. CNH wrote the manuscript. KSL participated in the writing of the manuscript and interpretation of results.

## Declaration of Interests

The authors declare no competing interests.

## Methods

### Cell Filtering

Raw counts expression matrices downloaded from GEO (accession IDs GSM1626793, GSM2743164) were filtered for high-quality cells prior to downstream analysis. The cumulative sum of total UMI counts for each cell was plotted along with the slope of the secant line to the curve as a function of rank-ordered cell. The distance between these two curves was used as a metric for determining the rate of diminishing cell quality. The cell number at which this distance was 50 % of its maximum was chosen as a cutoff, with cells contributing less UMI counts were removed. Next, a 100-component PCA and UMAP with n_neighbors value of 0.5 % of the total cells in the dataset were used to visualize cell populations and manually gate out clusters containing high mitochondrial counts, indicating dead cells.

Process shown in: github.com/KenLauLab/DR-structure-preservation/dev/QC.ipynb

**Figure 1.**
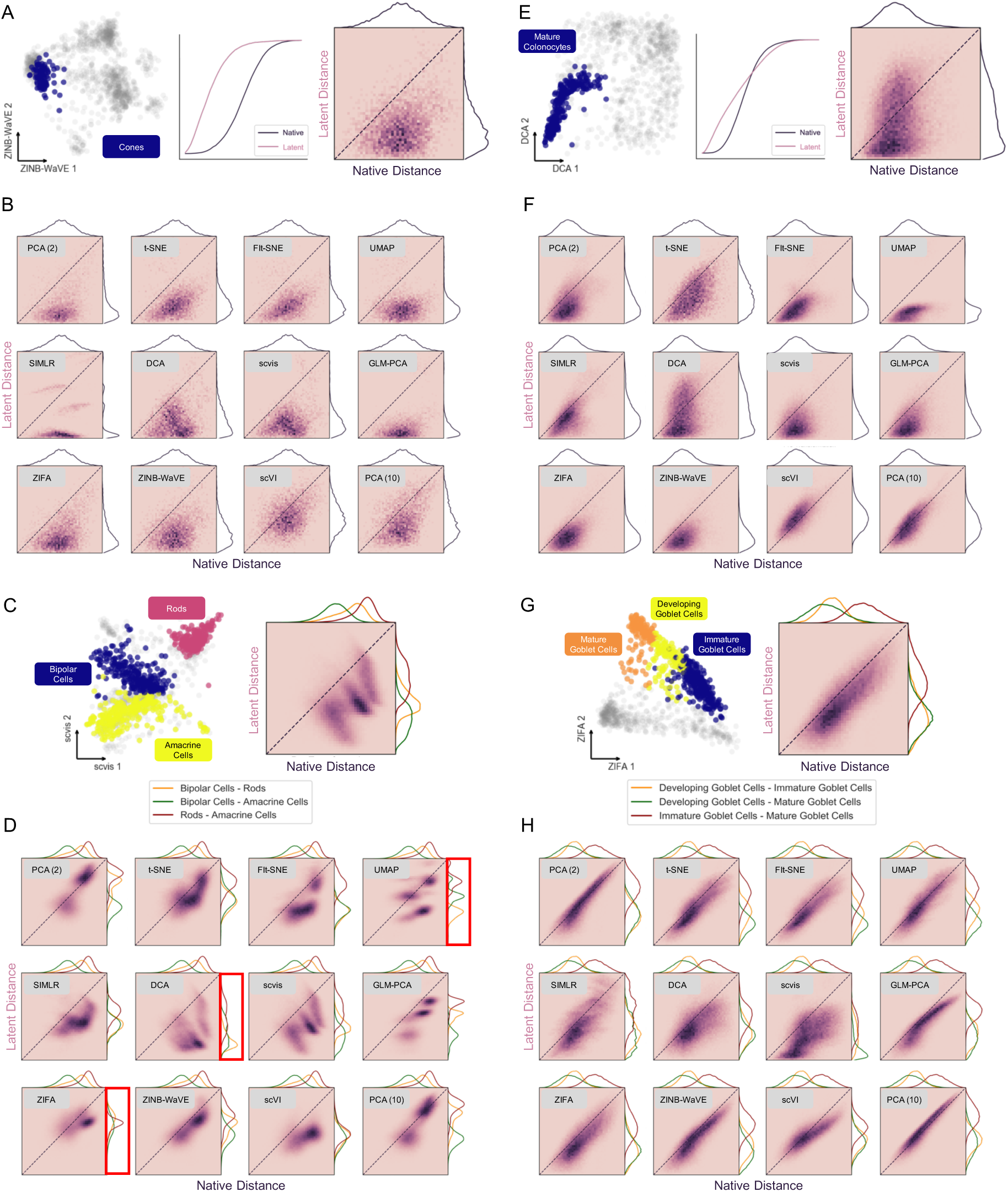
Local structure preservation analysis of 11 dimensionality reduction methods on discrete and continuous scRNA-seq datasets. A) Example 2D projection of mouse retina data using ZINB-WaVE and overlay of cone cells (left). Cumulative distance distributions within cones cluster for native and latent spaces (middle) and 2D histogram representing Pearson correlation between the two (right). B) 2D histograms of cell distance correlations within cone cell cluster for evaluated latent spaces. C) Same as in A for distances between bipolar, amacrine, and rod cell clusters, using scvis projection. D) Same as in B for distances between bipolar, amacrine, and rod cells. Methods that rearranged cluster ordering are highlighted in red. E) Example 2D projection of mouse colon data using DCA and overlay of mature colonocytes (left). Cumulative distance distributions within colonocyte cluster for native and latent spaces (middle) and 2D histogram representing Pearson correlation between the two (right). F) 2D histograms of cell distance correlations within mature colonocyte cluster for evaluated latent spaces. G) Same as in E for distances between three goblet cell clusters, using ZIFA projection. H) Same as in F for distances between three goblet cell clusters. See Methods.

### Clustering

PhenoGraph (Levine *et al*., 2015) was used to perform Louvain clustering on both datasets in Python. To create coarse, ground-truth clusters, the algorithm was run on 100 principal components of all genes in each dataset. For the retina data, 100 PCs of 20,478 genes explained 33.5 % of the variance in the dataset. For the colon data, 100 PCs of 25,505 genes explained 54.0 % of the variance. *k* values of 50 and 100 for generating the Knn graph to seed the Louvain algorithm for the retina and colon datasets, respectively, were chosen to provide coarse clustering of major cell types. Nine resulting clusters for the retina dataset and six resulting clusters in the colon dataset were analyzed by Seurat’s FindAllMarkers and DoHeatmap functions (Butler *et al*., 2018) to obtain visualizations of up- and down-regulated genes in each cluster (Figure 2A,D).

Process shown in:

- github.com/KenLauLab/DR-structure-preservation/dev/cluster_ID.ipynb
- github.com/KenLauLab/DR-structure-preservation/dev/consensus_clustering.r

### Dimensionality Reduction

All dimensionality reduction was performed on feature-selected data containing the most variable genes in each dataset. Genes were rank-ordered by variance using the Pandas (version 0.22.0) DataFrame.var function in Python, and the top 500 were chosen. Each dimensionality reduction technique was run “out-of-the-box” with default parameters on the feature-selected data. DCA, scvis, scVI, ZINB-WaVE and GLM-PCA take raw, unnormalized counts as input. Developers of ZIFA recommend a log2 transformation of counts, which we first normalized to the maximum UMI count within each cell. Arcsinh-transformed counts normalized to the maximum UMI count in each cell were used for all other methods (t-SNE, FIt-SNE, UMAP, SIMLR, PCA).

Process shown in:

- github.com/KenLauLab/DR-structure-preservation/dev/global_eval.ipynb
- github.com/KenLauLab/DR-structure-preservation/dev/Rmethods.Rmd
- github.com/KenLauLab/DR-structure-preservation/dev/scvis_out/README.md

### Distance Metric Calculations

Mantel test for correlation between symmetric Euclidean distance matrices (Figure 3C,D,G,H) was performed using the skbio.stats.distance.mantel function from the scikit-bio package (version 0.5.4). Pearson correlation was performed for local distance preservation analysis between clusters, as the resulting cell distance matrices are not symmetrical (Figure 3C,D,G,H). The scipy.stats.pearsonr function from the scipy package (version 1.1.0) was used. The scipy.stats.wasserstein_distance function from the scipy package (version 1.1.0) was used to calculated Earth Mover’s Distance between the flattened vectors containing unique distances between all cells in the dataset (upper triangle of distance matrix, Figure 3B,C,F,G), except for local comparisons between clusters, where the entire flattened matrix was used as the cell-cell distance matrices are not symmetrical (Figure 3C,D,G,H). A Knn graph with K = 30 was constructed using sklearn.neighbors.kneighbors_graph function from the scikit-learn package (version 0.20.0). Knn preservation was calculated as the percentage of elements in the Knn graph matrix that are conserved.

Functions used for above calculations can be found in:

- github.com/KenLauLab/DR-structure-preservation/dev/global_eval.ipynb
- github.com/KenLauLab/DR-structure-preservation/dev/local_eval.ipynb
- github.com/KenLauLab/DR-structure-preservation/dev/neighborhood_eval.ipynb

### Visualization

Cumulative cell distance distributions were plotted from the upper triangle of symmetrical cell distance matrices (using triu_indices function from the numpy Python package (version 1.16.3)). The histogram and cumsum functions numpy Python package (version 1.16.3) were used to plot cumulative distribution functions using n/100 bins, where n is the length of the flattened distance vector. Unique distance correlation was visualized using the JointGrid and kdeplot functions from the seaborn package (version 0.9.0), as well as the pyplot.hist2d function from the matplotlib package (version 3.0.1).

Functions used for above visualizations can be found in:

- github.com/KenLauLab/DR-structure-preservation/fcc_utils.py

### Lead Contact and Code Availability

- Further information and requests for resources and reagents should be directed to and will be fulfilled by the Lead Contact, KSL (ken.s.lau@ vanderbilt.edu).
- All code for this project is available at github.com/KenLauLab/DR-structure-preservation.
- Original data for this project is available on GEO:
  - Accession ID GSM1626793 (mouse retina, Macosko *et al*., 2015)
  - Accession ID GSM2743164 (mouse colon, Herring, Banerjee, *et al*., 2018)

## Supplemental Information

**Figure S1, Related to Figure 1.**
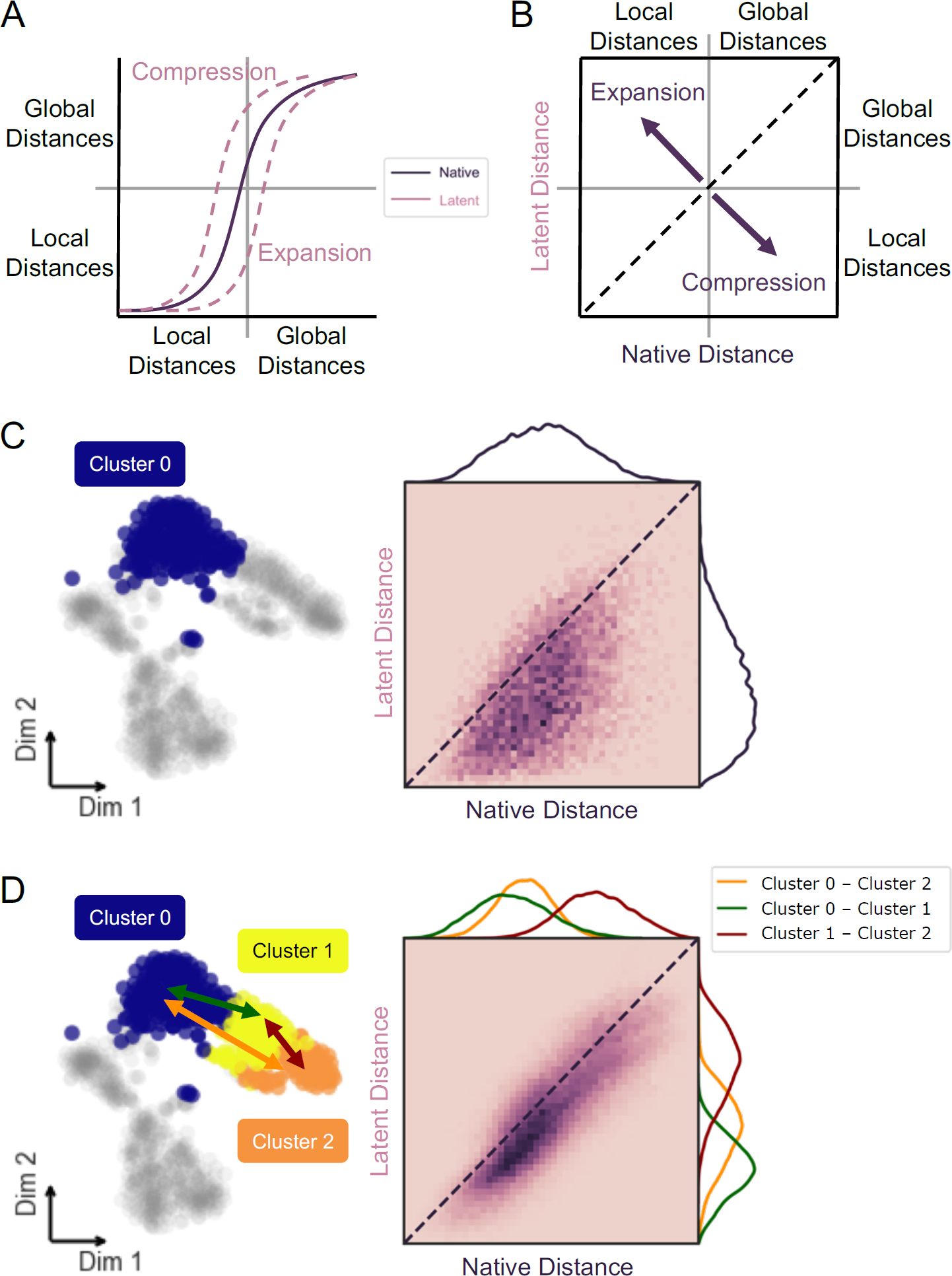
Interpretation of data structure preservation analysis. A) Small distances in cumulative distance distribution represent local cell similarity (within cluster), while large distances represent global relationships and arrangement of data (between clusters). A distribution shift left indicates compression of distances from native to latent space, while a shift right results from expansion or exaggeration of native distances. B) Correlation of latent to native distances; dispersion below identity line (dashed) indicates compression of distances from native to latent space, while dispersion above identity results from expansion of native distances in low-dimensional space. C) Substructure analysis uses same framework as Figure 1 on isolated subset of data to measure intra-cluster distance preservation and determine contribution to global structure. D) Distribution of distances from all cells in one cluster to another define relative substructure. Intercluster distances are measured pairwise to interrogate cluster arrangement in latent compared to native space.

**Figure S2, Related to Figure 2.**
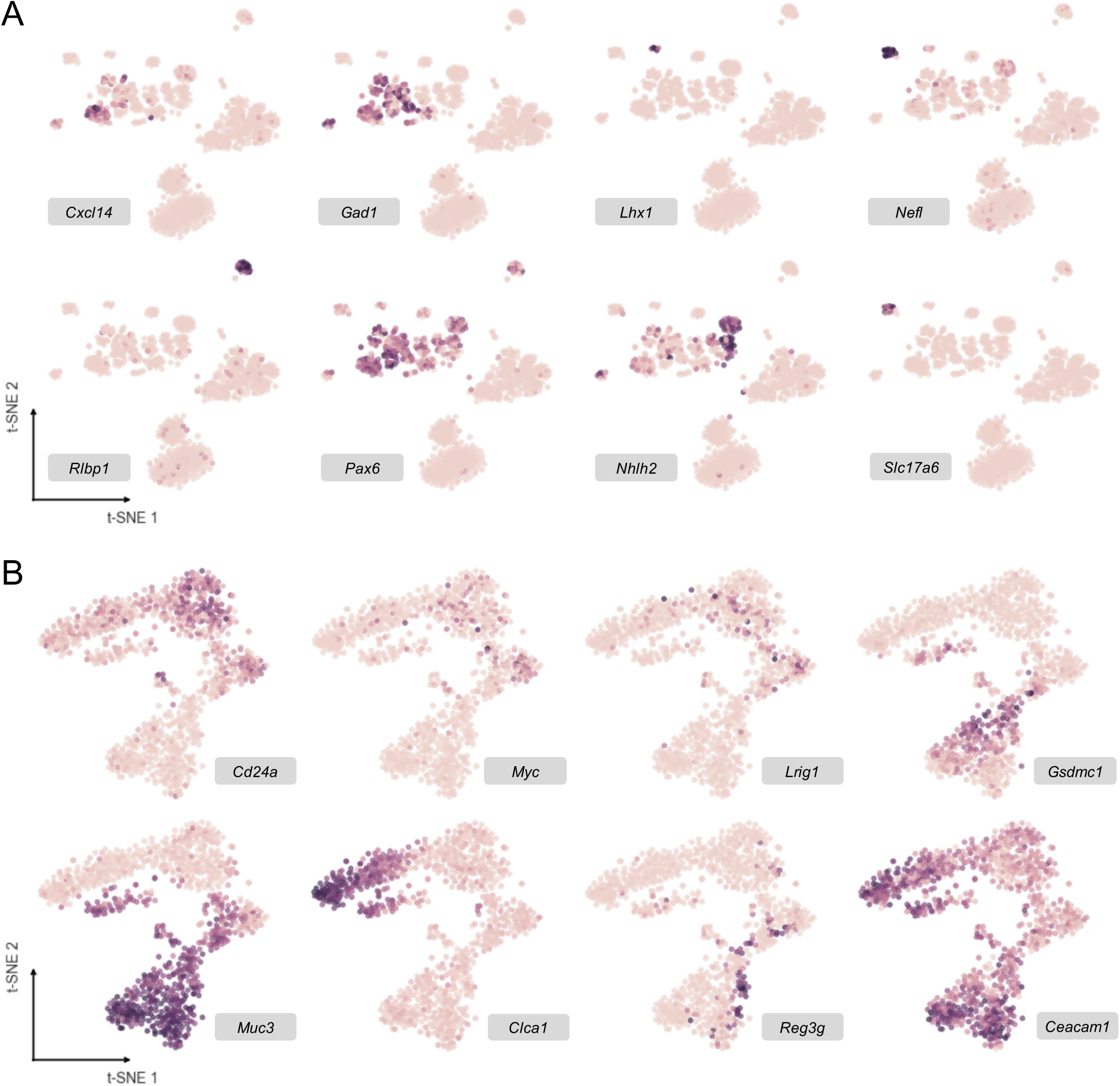
t-SNE visualizations from Figure 2 with overlay of arcsinh-normalized expression of marker genes. Used to assign cell type to Louvain clusters for retina (A) and colon (B) datasets.

**Figure S3, Related to Figure 3.**
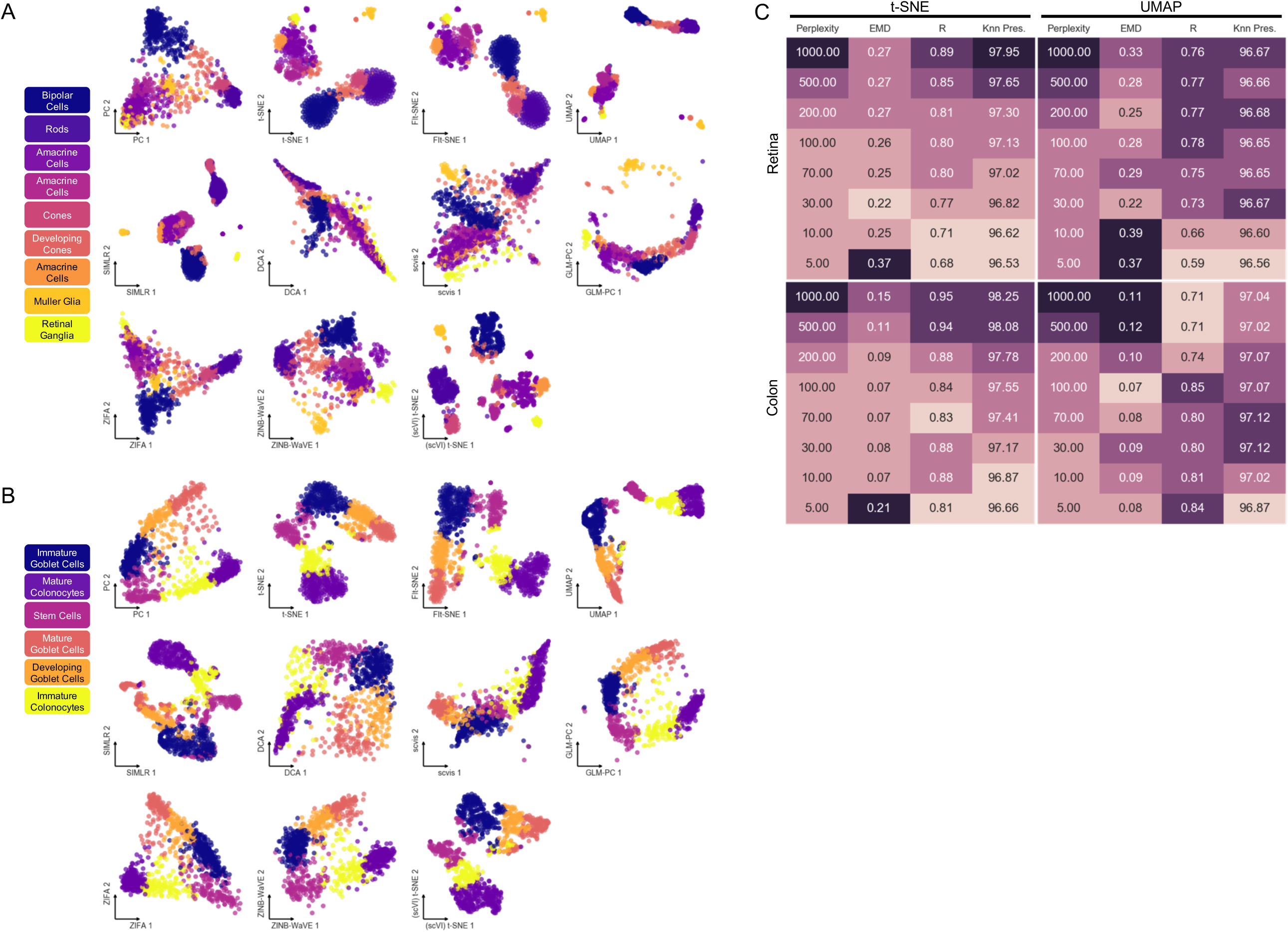
Low-dimensional projections from 11 evaluated dimensionality reduction methods with overlay of consensus Louvain clusters for retina (A) and colon (B) data. These embeddings were generated using the 500 most variable genes in each dataset. C) Resulting distance metrics from titration of perplexity parameter in t-SNE and UMAP (n_neighbors) on retina (discrete) and colon (continuous) datasets.

**Figure S4, Related to Figure 4.**
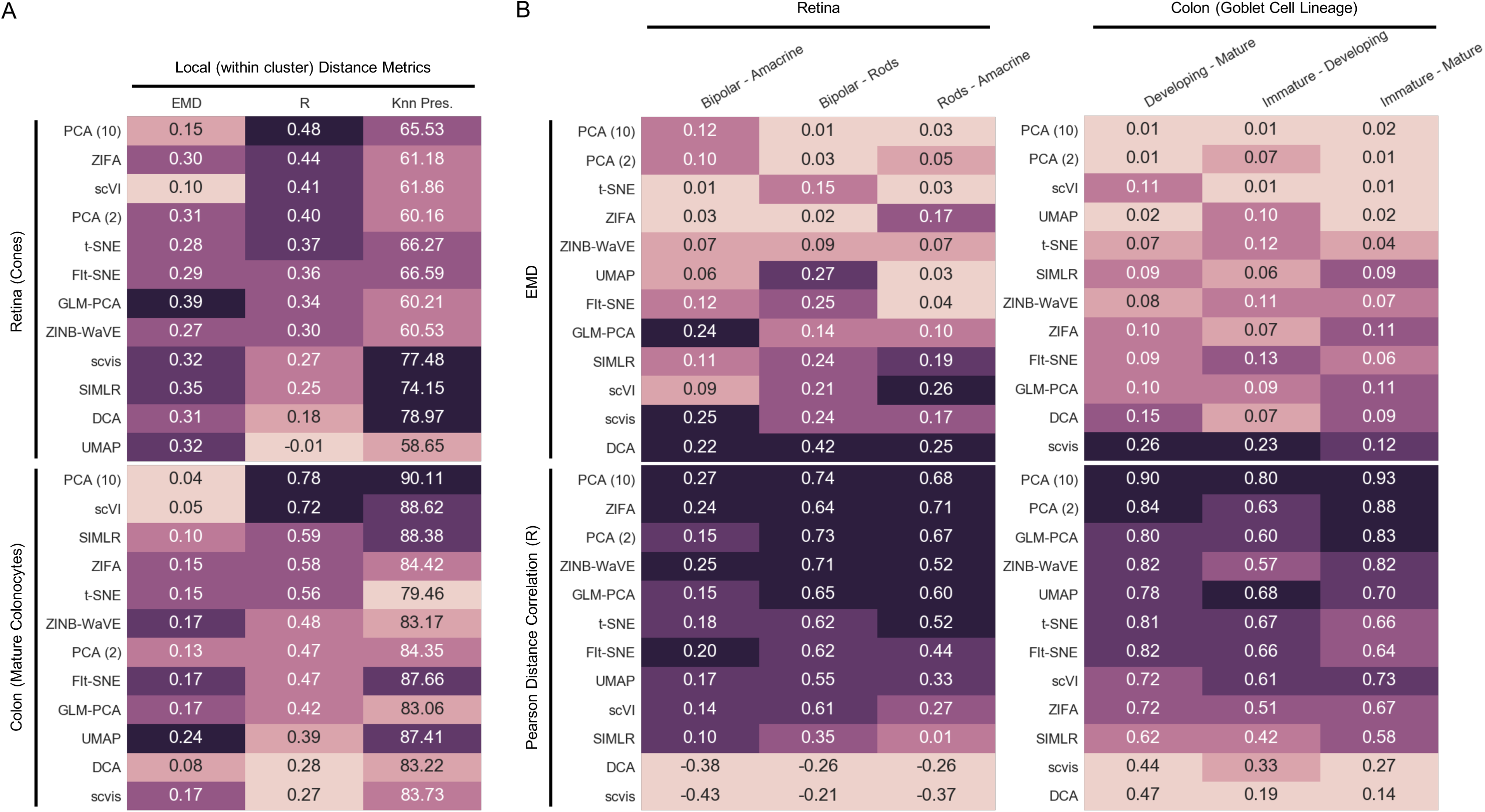
Summary of substructure analysis of discrete and continuous data. A) Local (within cluster) distance preservation metrics for cone cell cluster and mature colonocyte cluster from retina and colon datasets, respectively. B) EMD and distance correlation values for pairwise distance distributions between bipolar cells, rod cells, and amacrine cells in retina dataset, and three clusters along developing goblet cell lineage in colon dataset.

## Key Resources Table

| RESOURCE | SOURCE | IDENTIFIER |
| --- | --- | --- |
| Deposited Data |  |  |
| Mouse retina scRNA-seq data | Macosko *et al.*, 2015 | GEO: GSM1626793 |
| Mouse colon scRNA-seq data | Herring, Banerjee, *et al.*, 2018 | GEO: GSM2743164 |
| Software and Algorithms |  |  |
| Python version 3.6 | Python Software Foundation | https://www.python.org/downloads/ |
| R version 3.5 | The R Foundation | https://www.r-project.org |
| Seurat version 3.0.0 | Butler *et al.*, 2018 | https://satijalab.org/seurat/ |
| numpy version 1.16.3 | Oliphant, 2006 | http://www.numpy.org |
| scipy version 1.1.0 | Oliphant, 2007 | http://www.scipy.org/ |
| pandas version 0.22.0 | McKinney *et al.*, 2010 | https://pandas.pydata.org |
| scikit-learn version 0.20.0 | Pedregosa *et al.*, 2011 | https://scikit-learn.org |
| scikit-bio version 0.5.4 | Python Software Foundation | http://scikit-bio.org/ |
| PhenoGraph 1.5.2 | Levine *et al.*, 2015 | https://github.com/jacoblevine/PhenoGraph |
| umap-learn version 0.3.6 | McInnes and Healy, 2018 | https://github.com/lmcinnes/umap |
| scanpy version 1.3.2 | Wolf, Angerer and Theis, 2018 | https://github.com/theislab/scanpy |
| DCA version 0.2.2 | Eraslan *et al.*, 2019 | https://github.com/theislab/dca |
| scVI version 0.2.1 | Lopez *et al.*, 2018 | https://github.com/YosefLab/scVI |
| scvis version | Ding, Condon and Shah, 2018 | https://github.com/shahcompbio/scvis |
| SIMLR version 1.8.1 | Wang *et al.*, 2017 | https://github.com/BatzoglouLabSU/SIMLR |
| GLM-PCA | Townes *et al.*, 2019 | https://github.com/willtownes/scrna2019 |
| ZINB-WaVE version 1.4.2 | Risso *et al.*, 2018 | https://bioconductor.org/packages/zinbwave |
| ZIFA version 0.1 | Pierson and Yau, 2015 | https://github.com/epierson9/ZIFA |
| Flt-SNE | Linderman *et al.*, 2019 | https://github.com/KlugerLab/Flt-SNE |

## Notes

https://github.com/KenLauLab/DR-structure-preservation

## References

Becht, E. et al. (2018) ‘Dimensionality reduction for visualizing single-cell data using UMAP’, Nature Biotechnology. Nature Publishing Group. doi: 10.1038/nbt.4314.

Belkina, A. C. et al. (2018) ‘Automated optimal parameters for T-distributed stochastic neighbor embedding improve visualization and allow analysis of large datasets’, bioRxiv, p. 451690. doi: 10.1101/451690.

Van den Berge, K. et al. (2019) ‘Trajectory-based differential expression analysis for single-cell sequencing data’, bioRxiv. Cold Spring Harbor Laboratory, p. 623397. doi: 10.1101/623397.

Butler, A. et al. (2018) ‘Integrating single-cell transcriptomic data across different conditions, technologies, and species’, Nature Biotechnology. Nature Publishing Group, 36(5), pp. 411–420. doi: 10.1038/nbt.4096.

Chen, B., Herring, C. A. and Lau, K. S. (2018) ‘pyNVR: investigating factors affecting feature selection from scRNA-seq data for lineage reconstruction’, Bioinformatics. Edited by I. Birol. doi: 10.1093/bioinformatics/bty950.

Ding, J., Condon, A. and Shah, S. P. (2018) ‘Interpretable dimensionality reduction of single cell transcriptome data with deep generative models’, Nature Communications. Nature Publishing Group, 9(1), p. 2002. doi: 10.1038/s41467-018-04368-5.

Eraslan, G. et al. (2019) ‘Single-cell RNA-seq denoising using a deep count autoencoder’, Nature Communications. Nature Publishing Group, 10(1), p. 390. doi: 10.1038/s41467-018-07931-2.

Herring, C. A., Chen, B., et al. (2018) ‘Single-Cell Computational Strategies for Lineage Reconstruction in Tissue Systems’, Cellular and Molecular Gastroenterology and Hepatology. Elsevier, 5(4), pp. 539–548. doi: 10.1016/J.JCMGH.2018.01.023.

Herring, C. A., Banerjee, A., et al. (2018) ‘Unsupervised Trajectory Analysis of Single-Cell RNA-Seq and Imaging Data Reveals Alternative Tuft Cell Origins in the Gut’, Cell Systems. Cell Press, 6(1), pp. 37–51.e9. doi: 10.1016/J.CELS.2017.10.012.

Kiselev, V. Y. et al. (2017) ‘SC3: consensus clustering of single-cell RNA-seq data’, Nature Methods. Nature Publishing Group, 14(5), pp. 483–486. doi: 10.1038/nmeth.4236.

Klein, A. M. et al. (2015) ‘Droplet barcoding for single-cell transcriptomics applied to embryonic stem cells.’, Cell. NIH Public Access, 161(5), pp. 1187–1201. doi: 10.1016/j.cell.2015.04.044.

Kobak, D. and Berens, P. (2019) ‘The art of using t-SNE for single-cell transcriptomics’, bioRxiv. Cold Spring Harbor Laboratory, p. 453449. doi: 10.1101/453449.

Larsson, E. et al. (2012) ‘Analysis of gut microbial regulation of host gene expression along the length of the gut and regulation of gut microbial ecology through MyD88.’, Gut. BMJ Publishing Group, 61(8), pp. 1124–31. doi: 10.1136/gutjnl-2011-301104.

Lepourcelet, M. et al. (2005) ‘Insights into developmental mechanisms and cancers in the mammalian intestine derived from serial analysis of gene expression and study of the hepatoma-derived growth factor (HDGF)’, Development. The Company of Biologists Ltd, 121(10), pp. 3163–3174. doi: 10.1242/dev.01579.

Levina, E. and Bickel, P. (2001) ‘The Earth Mover’s distance is the Mallows distance: some insights from statistics’, in Proceedings Eighth IEEE International Conference on Computer Vision. ICCV 2001. IEEE Comput.Soc, pp. 251–256. doi: 10.1109/ICCV.2001.937632.

Levine, J. H. et al. (2015) ‘Data-Driven Phenotypic Dissection of AML Reveals Progenitor-like Cells that Correlate with Prognosis.’, Cell. Elsevier, 162(1), pp. 184–97. doi: 10.1016/j.cell.2015.05.047.

Linderman, G. C. et al. (2017) ‘Efficient Algorithms for t-distributed Stochastic Neighborhood Embedding’. doi: 10.1038/s41592-018-0308-4.

Linderman, G. C. et al. (2019) ‘Fast interpolation-based t-SNE for improved visualization of single-cell RNA-seq data’, Nature Methods. Nature Publishing Group, 16(3), pp. 243–245. doi: 10.1038/s41592-018-0308-4.

Lopez, R. et al. (2018) ‘Deep generative modeling for single-cell transcriptomics’, Nature Methods. Nature Publishing Group, 15(12), pp. 1053–1058. doi: 10.1038/s41592-018-0229-2.

Van der Maaten, L. and Hinton, G. (2008) ‘Visualizing Data using t-SNE’, Journal of Machine Learning Research, 9(Nov), pp. 2579–2605. Available at: http://www.jmlr.org/papers/v9/vandermaaten08a.html (Accessed: 11 May 2019).

Macosko, E. Z. et al. (2015) ‘Highly Parallel Genome-wide Expression Profiling of Individual Cells Using Nanoliter Droplets’, Cell. Cell Press, 161(5), pp. 1202–1214. doi: 10.1016/J.CELL.2015.05.002.

Mantel, N. (1967) ‘The detection of disease clustering and a generalized regression approach.’, Cancer research. American Association for Cancer Research, 27(2), pp. 209–20. Available at: http://www.ncbi.nlm.nih.gov/pubmed/6018555 (Accessed: 12 June 2019).

Mcinnes, L. and Healy, J. (2018) UMAP: Uniform Manifold Approximation and Projection for Dimension Reduction. Available at: https://arxiv.org/pdf/1802.03426.pdf (Accessed: 29 October 2018).

McKinney, W. et al. (2010) ‘Data structures for statistical computing in python. In Proceedings of the 9th Python in Science Conference. pp. 51–56.

Oliphant, T. E. (2006) ‘A guide to NumPy’, Trelgol Publishing USA.

Oliphant, T. E. (2007) ‘Python for Scientific Computing’, Computing in Science & Engineering. IEEE Computer Society, 9(3), pp. 10–20. doi: 10.1109/MCSE.2007.58.

Pedregosa, F. et al. (2011) ‘Scikit-learn: Machine Learning in Python’, Journal of Machine Learning Research, 12(Oct), pp. 2825–2830. Available at: http://jmlr.csail.mit.edu/papers/v12/pedregosa11a.html (Accessed: 17 June 2019).

Pierson, E. and Yau, C. (2015) ‘ZIFA: Dimensionality reduction for zero-inflated single-cell gene expression analysis’, Genome Biology. BioMed Central, 16(1), p. 241. doi: 10.1186/s13059-015-0805-z.

Qiu, X. et al. (2017) ‘Reversed graph embedding resolves complex single-cell trajectories’, Nature Methods. Nature Publishing Group, 14(10), pp. 979–982. doi: 10.1038/nmeth.4402.

Regev, A. et al. (2017) ‘The Human Cell Atlas’, eLife, 6. doi: 10.7554/eLife.27041.

Rheaume, B. A. et al. (2018) ‘Single cell transcriptome profiling of retinal ganglion cells identifies cellular subtypes’, Nature Communications. Nature Publishing Group, 9(1), p. 2759. doi: 10.1038/s41467-018-05134-3.

Risso, D. et al. (2018) ‘A general and flexible method for signal extraction from single-cell RNA-seq data’, Nature Communications. Nature Publishing Group, 9(1), p. 284. doi: 10.1038/s41467-017-02554-5.

Rubner, Y., Tomasi, C. and Guibas, L. J. (1998) ‘A metric for distributions with applications to image databases’, in Sixth International Conference on Computer Vision (IEEE Cat. No.98CH36271). Narosa Publishing House, pp. 59–66. doi: 10.1109/ICCV.1998.710701.

Rubner, Y., Tomasi, C. and Guibas, L. J. (2000) ‘The Earth Mover’s Distance as a Metric for Image Retrieval’, International Journal of Computer Vision. Kluwer Academic Publishers, 40(2), pp. 99–121. doi: 10.1023/A:1026543900054.

Saelens, W. et al. (2019) ‘A comparison of single-cell trajectory inference methods’, Nature Biotechnology. Nature Publishing Group, 37(5), pp. 547–554. doi: 10.1038/s41587-019-0071-9.

Sorzano, C. O. S., Vargas, J. and Montano, A. P. (2014) ‘A survey of dimensionality reduction techniques’. Available at: http://arxiv.org/abs/1403.2877 (Accessed: 13 June 2019).

Tamura, M. et al. (2007) ‘Members of a novel gene family, Gsdm, are expressed exclusively in the epithelium of the skin and gastrointestinal tract in a highly tissue-specific manner’, Genomics. Academic Press, 89(5), pp. 618–629. doi: 10.1016/J.YGENO.2007.01.003.

Townes, F. W. et al. (2019) ‘Feature Selection and Dimension Reduction for Single Cell RNA-Seq based on a Multinomial Model’, bioRxiv. Cold Spring Harbor Laboratory, p. 574574. doi: 10.1101/574574.

Tsuyuzaki, K. et al. (2019) ‘Benchmarking principal component analysis for large-scale single-cell RNA-sequencing’, bioRxiv. Cold Spring Harbor Laboratory, p. 642595. doi: 10.1101/642595.

Tusi, B. K. et al. (2018) ‘Emergence of the erythroid lineage from multipotent hematopoiesis’, bioRxiv. Cold Spring Harbor Laboratory, p. 261941. doi: 10.1101/261941.

Wagner, D. E. et al. (2019) Single-cell mapping of gene expression landscapes and lineage in the zebrafish embryo Downloaded from. Available at: http://science.sciencemag.org/ (Accessed: 18 February 2019).

Wang, B. et al. (2017) ‘Visualization and analysis of single-cell RNA-seq data by kernel-based similarity learning’, Nature Methods. Nature Publishing Group, 14(4), pp. 414–416. doi: 10.1038/nmeth.4207.

Werman, M., Peleg, S. and Rosenfeld, A. (1985) ‘A distance metric for multidimensional histograms’, Computer Vision, Graphics, and Image Processing. Academic Press, 32(3), pp. 328–336. doi: 10.1016/0734-189X(85)90055-6.

Wolf, F. A., Angerer, P. and Theis, F. J. (2018) ‘SCANPY: large-scale single-cell gene expression data analysis’, Genome Biology. BioMed Central, 19(1), p. 15. doi: 10.1186/s13059-017-1382-0.

Zeisel, A. et al. (2015) ‘Brain structure. Cell types in the mouse cortex and hippocampus revealed by single-cell RNA-seq.’, Science (New York, N.Y.). American Association for the Advancement of Science, 347(6226), pp. 1138–42. doi: 10.1126/science.aaa1934.

